# Immediate impact of transcranial magnetic stimulation on brain structure: short-term neuroplasticity following one session of cTBS

**DOI:** 10.1101/2020.11.03.366534

**Authors:** JeYoung Jung, Matthew A. Lambon Ralph

**Affiliations:** School of Psychology, University of Nottingham, UK; Precision Imaging Beacon of Excellence, University of Nottingham, UK; MRC Cognition and Brain Science Unit (CBU), University of Cambridge, UK

**Keywords:** Structural plasticity, theta burst stimulation, voxel-based morphometry, anterior temporal lobe, cTBS combined fMRI

## Abstract

Recent evidence demonstrates that activation-dependent neuroplasticity on a structural level can occur in a short time (2 hour or less) in humans. However, the exact time scale of structural plasticity in adult human brain remains unclear. Using voxel-based morphometry (VBM), we investigated changes in gray matter (GM) after one session of continuous theta-burst stimulation (cTBS) delivered to the anterior temporal lobe (ATL). Twenty-five participants were received cTBS over the left ATL or occipital pole as a control site outside of the scanner and had structural and functional imaging. During functional imaging, participants performed a semantic association task. VBM result revealed decreased GM in the left ATL and right cerebellum after ATL stimulation compared to the control stimulation. In addition, cTBS over the left ATL induced slower reaction time in sematic task performance, reduced regional activity at the target site, and altered functional connectivity between the left and right ATL during semantic processing. Furthermore, the ATL GM changes were associated with the functional connectivity changes in the ATL-connectivity during semantic processing. These structural alterations are mirrored by functional changes in cortical excitability attributed to the GM changes and demonstrate the rapid dynamics of cortical plasticity. Our findings support fast adjusting neuronal systems, such as postsynaptic morphology changes and neuronal turnover. Our results suggest that TBS is able to produce powerful changes in regional synaptic activity in human adult brain.

## 1. Introduction

Neuroplasticity refers to the brain’s ability to reorganize itself in response to environmental changes and involves a complex, multilayer process including the molecular, synaptic, electrophysiological and structural organization level. The scope of neuroplasticity encompasses functional forms including short-term weakening and strengthening of existing synapses through long-term potentiation (LTP) and long-term depression (LTD) and structural types such as synaptogenesis, gliogenesis, and neurogenesis (Bruel-Jungerman et al., 2007; Butz et al., 2009; Holtmaat and Svoboda, 2009). Although traditional neuroscience research has focused on functional forms of neuroplasticity in the adult brain, structural types of plasticity play a critical role in adaptation to environmental changes and diseases such as stroke (Pekna et al., 2012).

Structural neuroplasticity are thought to be more slow and infrequent than the functional plasticity (Bruel-Jungerman et al., 2007). However, the timescale of structural plasticity is still poorly understood. Animal studies have showed that neurogenesis occurs within days, whereas local morphological changes such as formation of new synapses and dendrites can arise on shorter time period, less than a day (Bruel-Jungerman et al., 2007; Butz et al., 2009; Holtmaat and Svoboda, 2009; Trachtenberg et al., 2002). A recent study showed that regional structural changes (increased dendrite spines) in rodent’s brain happened immediately after a short-term training (about 1 hour) (Xu et al., 2009). In human, a long period of training (weeks or months) induces such structural changes (Draganski et al., 2004; Zatorre et al., 2012). In human adults, Sagi et al (2012) demonstrated that only 2 hour of learning resulted in brain structural alterations using diffusion tensor imaging. Participants in the learning group performed a spatial learning and memory task for 90min on average and microstructural changes of hippocampus and parahippocampus were observed with improved task performance in the learning group compared to the control group (no learning). Their findings were replicated in a subsequent rat study such that there was structural remodelling of the rat hippocampus following 2 hour of water maze task (Sagi et al., 2012). These studies provide evidence that structural plasticity can occur short time scales (∼ hours) in adults. It is important to understand what extent the structural plasticity arises in adult’s brain following environment demands and disease because this type of cortical plasticity is associated with short-and long-term therapeutic effects.

Theta-burst transcranial magnetic stimulation (TBS) as an effective repetitive transcranial magnetic stimulation (rTMS) protocol modulates cortical excitability with a short period of stimulation through changes in synaptic strength (e.g., LTP and LTD) (Huang et al., 2005). TBS has increasingly and successfully been used to explore the mechanisms and consequences of functional plasticity in human cortex (Agnew et al., 2018; Hartwigsen et al., 2013; Jung and Lambon Ralph, 2016; Valchev et al., 2016). Animal studies demonstrated that TBS can induce immediate and prolonged functional and structural plasticity (For a review, see Funke and Benali, 2011). TBS modulated the GABA-synthesizing enzymes, presynaptic GABA transporters, and cortical inhibitory interneurons (Funke and Benali, 2010; Trippe et al., 2009). Specifically, continuous TBS (cTBS) reduced the number of calbindin expressing interneuron, whereas intermittent TBS (iTBS) decreased in parvalbumin expressing cells right after the stimulation (Benali et al., 2011). In human, TBS induced morphological changes were observed in gray matter (GM) and white matter (WM) (Allendorfer et al., 2012; May et al., 2007). May et al (2007) used voxel-based morphometry (VBM) to detect structural alterations following 5 days 1Hz rTMS over the left superior temporal gyrus. They reported increased GM in the targeted area as well as a transient increase and decrease of GM in the contralateral region.

Previously, we showed that an inhibitory, cTBS over the anterior temporal lobe (ATL) induced rapid, adaptive functional reorganization in the semantic representation system, revealing regional activity changes at the target region and connected homologues region as well as altered functional connectivity between them (Jung and Lambon Ralph, 2016). Here, we re-analysed the data set from our previous study with respect to structural plasticity induced by cTBS, a question that we did not address in the original publication. Healthy subjects received cTBS applied over left ATL and a control stimulation over the occipital pole (Oz) (Jung and Lambon Ralph, 2016). After the stimulation, structural images were obtained and analysed using VBM to assess changes in GM and WM. We hypothesized that cTBS over the ATL would decrease the brain morphology at the target region as well as the related white matter tract. In addition, we predicted that the structural alteration induced by cTBS would be associated with short-term functional neuroplasticity in the semantic representation system found in fMRI.

## 2. Materials and Methods

### 2.1 Subjects

All data have previously been included in a publication in a cTBS-fMRI study (Jung and Lambon Ralph, 2016). We re-analysed the fMRI data set with respect to microstructural changes in gray and white matter caused by cTBS. Accordingly, data from 25 healthy, right-handed native English speakers were included (7 males, mean age, 21.9 ± 3.7 years, range from 19 to 34 years). Data from 2 subjects were excluded in the analysis due to excessive head movements during fMRI (over a voxel). Handedness was assessed using the Edinburgh Handedness Inventory (Oldfield, 1971). All subjects provided informed written consent. The study was approved by the local ethics committee.

### 2.2 Experimental design and procedure

A detailed description of the procedure has been previously published (Jung and Lambon Ralph, 2016). We here summarized the important steps. We used a within-subject design to test the effects of one-session cTBS on gray and white matter density. Each subject participated in two MRI sessions (Fig. 1). In each session, cTBS was applied prior to the MRI outside of the scanner: the left ATL stimulation and Oz stimulation as a control site. Sessions were separated at least one week to avoid carry-over effects. The order of ATL and Oz cTBS was counterbalanced between subjects.

**Figure 1.**
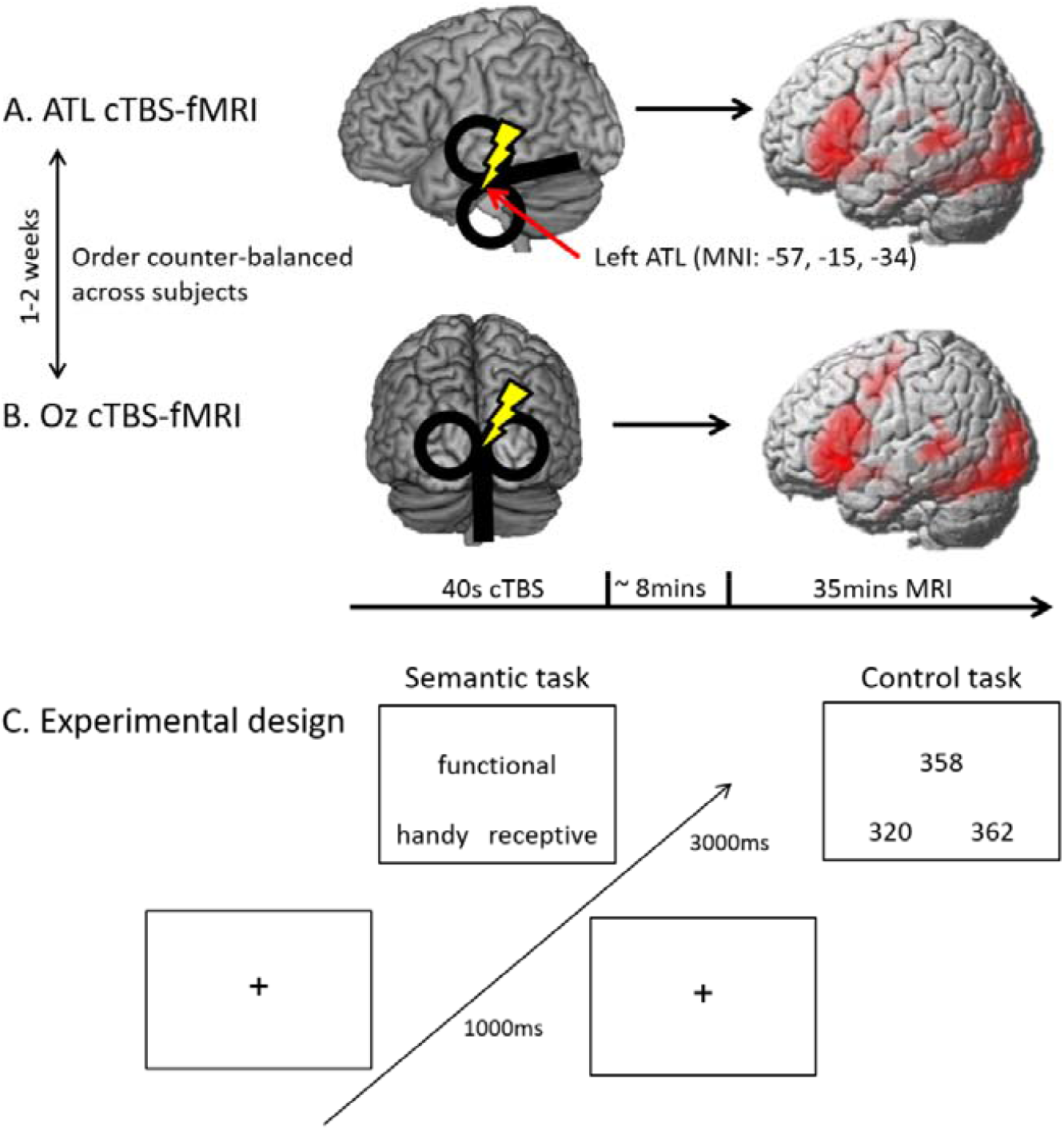
Experimental design and procedures. A) cTBS over the left ATL and following fMRI with tasks. B) cTBS over the occipital pole as a control stimulation. C) Experimental design

In the MRI, all subject performed a semantic judgement task and a number judgement task as a control task (Fig. 1C). In the semantic task, subjects saw 3 words on the screen and select which of 2 words (bottom) was more related to a target word (top) in meaning. In the control task, subjects saw 3 numbers and chose which of 2 numbers (bottom) was closer to the target number (target) in numerical value. Each trial started with 1s fixation followed by the stimuli presented for a fixed duration of 3s. A block design fMRI was used with 3 condition blocks: semantic, control and fixation. Each task block had 4 trials of a task and the fixation blocks (8s) were interleaved between task blocks. E-prime software (Psychology Software Tools Inc. Pittsburgh, PA, USA) was used to display stimuli and to record responses.

### 2.3 Theta-burst stimulation

cTBS (600 pulses at 50Hz for 40s) was delivered over the stimulation sites using a Magstim SuperRapid 2 with a figure-of-eight coil (70mm standard coil, MagStim Company, Whitland, UK) according to Huang et al (Huang et al., 2005). cTBS was applied during the ATL and Oz at 80% of the resting motor threshold (RMT). The mean of stimulation intensity was 49.1% ranging from 36% to 61%.

All subjects were scanned to obtain a high-resolution T1-weighted anatomical image (in-plane resolution of 1mm and a slice thickness of 1.8mm with an acquisition matrix 256 × 256 voxels) using a 3T Philips MR Achieva scanner prior to the experiment.

The target site was selected the coordinate for the ATL based on a previous distortion-corrected fMRI study (Visser et al., 2012) [MNI: -57, -15, -34]. The ATL coordinate was transformed into each subject’s native space by normalizing each subject’s MRI scan against the MNI template using Statistical Parametric Mapping software (SPM8, Wellcome Trust Centre for Neuroimaging, London, UK). Then, the inverse of each resulting transformation was used to convert the target MNI coordinate to the untransformed individual naïve space coordinate. These ATL coordinates were used to guide the frameless stereotaxy, Brainsight TMS-MRI co-registration system (Rogue Research, Montreal, Canada). The control site was the occipital pole (Oz) localised by international 10-20 system.

### 2.4 Image acquisition

MRI was performed on a 3T Philips Achieva scanner using an 8-element SENSE head coil with a SENCE factor 2.5. A high-resolution T1-weighted image was acquired using a 3D MPRAGE pulse sequence with 200 slices, in plane resolution 0.94 × 0.94 mm, slice thickness 0.9 mm, TR = 8.4 ms, TE = 3.9 ms. For fMRI, a dual-echo protocol developed by Halai et al (Halai et al., 2014) was used in order to compensate the signal dropout around rostral temporal areas (42 slices, 96 × 96 matric, 240 × 240 × 126 mm FOV, in-plane resolution 2.5 × 2.5 mm, slice thickness 3 mm, TR = 2.8 s, TE = 12 ms and 35 ms).

### 2.5 Voxel-based morphometry

Voxel-based morphometry (VBM) was performed in order to investigate the GM and WM changes using a VBM8 toolbox (http://dbm,neuro.uni-jena.de/vbm8) in SPM8. Preprocessing of the data involved normalization, segmentation, modulation, and smoothing (Ashburner and Friston, 2000). We created a customized GM and WM templates from subjects in this study (Oz stimulation secession). Normalization parameters were estimated using an optimized protocol in order to facilitate optimal segmentation by normalizing extracted GM and WM images to the customized GM and WM templates (Good et al., 2001). Then the optimized parameters were reapplied to the original brain images. The images were aligned with the MNI space, corrected for non-uniformities in signal intensity and partitioned into GM, WM, cerebrospinal fluid (CSF), and background. Modulation was performed for volume change correction by modulating each voxel with the Jacobian determinants derived from the spatial normalization, which allows us to test regional differences in the absolute amount of GM and WM (Ashburner and Friston, 2000). Finally all images were smoothed by convolving them with an isotropic Gaussian kernel of 8 mm full-width at half maximum.

Voxel-by-voxel statistical procedures using the GLM were performed to examine regional specific GM and WM changes between ATL and Oz stimulation. The random effect model performed paired t-tests for each subject’s scans using TMS sites (ATL vs. Oz), accounting for the total intracranial volume. The statistical threshold set for all contrasts was p < 0.001 uncorrected with an extent threshold of 100 contiguous voxels. We hypothesized that cTBS over the left ATL would alter the brain morphology in this region and related white matter tract based on the previous findings (Jung and Lambon Ralph, 2016). A priori region was defined as an 8mm sphere in the left ventral ATL (vATL MNI: -33, -9 -39). For the white matter tract, we used the anterior commissure map from NatBrainLab (http://www.natbrainlab.co.uk/atlas-maps). We applied a threshold of FWE-corrected p < 0.05 for the multiple comparisons correction after a small volume correction (SVC), using a priori region.

### 2.6 fMRI analysis

A detailed description of the fMRI analysis has been previously published (Jung and Lambon Ralph, 2016). We here summarized the important steps. The dual gradient echo images were extracted and combined using in-house Matlab (Halai et al., 2014). The combined images were realigned, coregistered, normalized to the structural image and smoothed with an 8 mm full-width half-maximum Gaussian filter using SPM8. General linear model (GLM) analysis was performed to set up a fixed-effect model with each task condition (semantic and control) and to assess differences in activation between the contrasts (semantic > control) using a random-effect model. Region of interest (ROI) analysis was performed using a priori ROI (vATL).

Dynamic causal modelling (DCM) was used to estimate effective connectivity between the bilateral ventral ATL after suppression of the left ATL by cTBS. DCM is to estimate and make directional inferences in a predefined set of brain regions in different experimental context (Friston et al., 2003). The DCM models were based on bilateral ATLs which showed significant TMS effects in previous analysis(Jung and Lambon Ralph, 2016). Thus the model consisted of bilateral, intrinsic connections between the ventral ATLs. Then we set up 3 possible modulatory connections between them, reflecting the changes in the intrinsic connections induced by the experimental conditions (ATL stimulation in this study). Bayesian model selection (Stephan et al., 2009) was applied to determine which DCM models were the most likely given the observed our fMRI data. The result showed the winning model having a connection from the left to right ATL during semantic processing. We extracted individual specific parameters of the winning model, including the intrinsic connectivity (left ventral ATL → right ventral ATL and right ventral ATL → left ventral ATL) and the modulatory connectivity (left ventral ATL → right ventral ATL) and explored the relationship between the connectivity and brain morphological changes induced by cTBS (Pearson’s correlation, p < 0.05).

## 3. Results

A detailed description of the results has been previously published (Jung and Lambon Ralph, 2016). We here summarized the key findings from the previous study and reported findings related to changes in GM and WM induced by cTBS. Behavioural results demonstrated the inhibitory cTBS effects (slower reaction time; RT, t(22) = −3.47, p < 0.005) after the ATL stimulation compared to the control stimulation during the semantic task (Fig. 2A). In fMRI results, cTBS over the left ATL induced the decreased task-induced activation in the ATL (t(22) = 3.79, p < 0.01) as well as a compensatory up-regulation in the homologues right ATL (t(22) = −2.32, p = 0.05) (Fig. 2B). Furthermore, the effective connectivity between the ATLs was modulated by cTBS, showing the compensatory facilitation from the right ATL (intact region) to the left ATL (lesioned region) and increased task-specific connectivity during semantic processing (left ATL → right ATL) (Fig. 2C, see the summary in Table S1). These results demonstrated the fast adaptive functional reorganization of semantic system after one session cTBS intervention.

**Figure 2.**
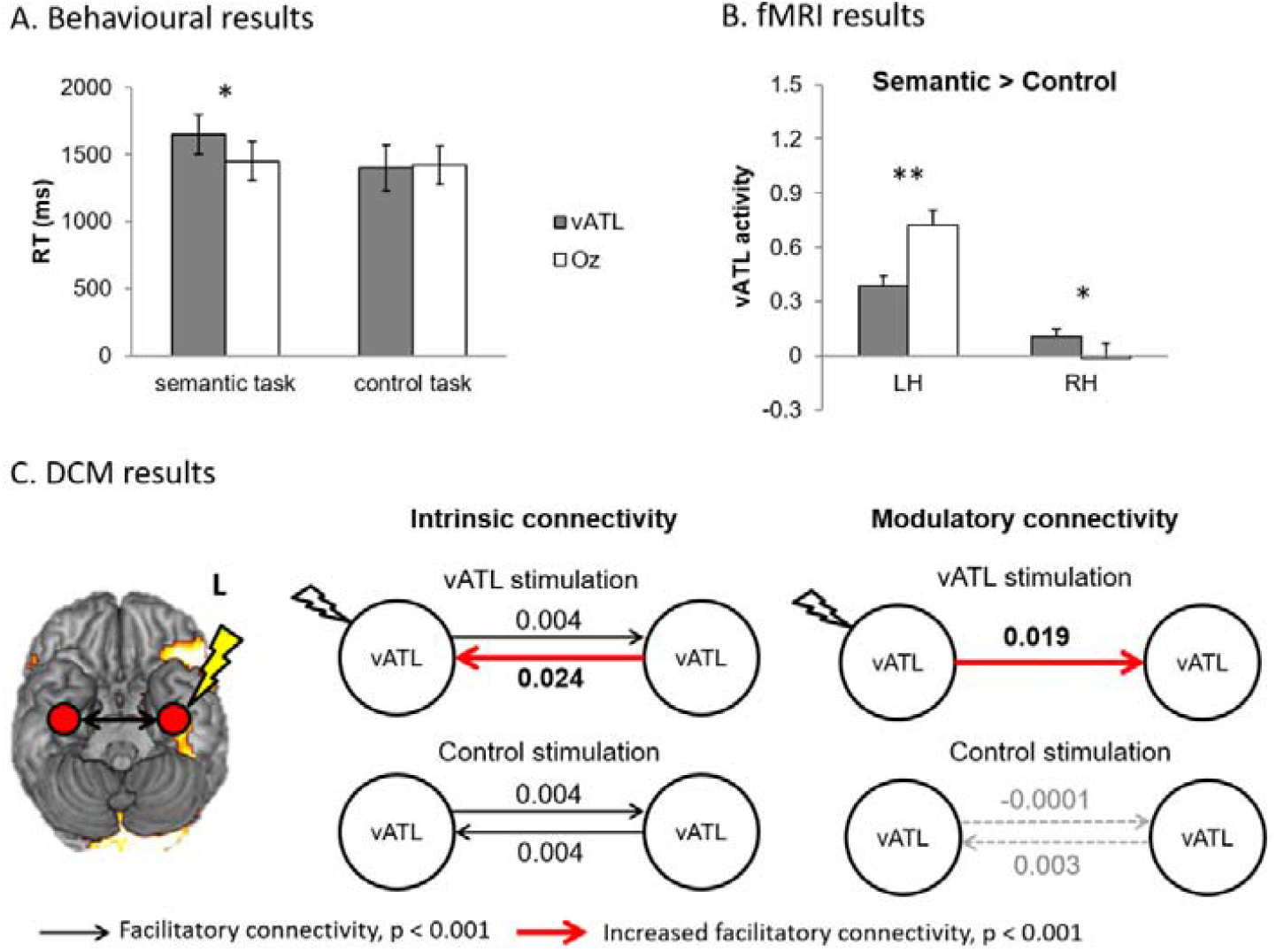
A) Behavioural results B) fMRI results – regional activity in the ventral ATL during semantic processing. Grey bar indicates the vATL stimulation. White bar represents the control stimulation. C) Dynamic causal modelling results. Black arrow represents significant facilitatory connectivity. Red arrow indicates increased facilitatory connectivity after ATL stimulation compared to the control stimulation. Dotted arrow represent non-significant connectivity.

VBM results showed that there was a significant transient decrease in the left ventral ATL (MNI: -36 -12 -24, cluster size = 108, p _SVC-FWE corrected_ = 0.005) after the ATL stimulation compared to the control stimulation (Fig. 3A). We also found a significant GM decrease in the right cerebellum (MNI: 17, -37, -48, cluster size = 1269, p _FWE-corrected_ < 0.001). There was no GM changes in the comparison of ATL stimulation > Oz stimulation. The dynamic pattern of the GM changes was specific to the ATL stimulation. Then, we investigated the relationship between the GM changes and functional short-term plasticity in the semantic system - effective connectivity between the ATLs. We found that there was a significant positive correlation between the ATL GM changes and ATL-connectivity. Participants with stronger connectivity between the left and right ATL showed greater GM in the left ventral ATL (intrinsic connectivity: r = 0.44, p = 0.017, modulatory connectivity: r = 0.47, p = 0.012) (Fig. 3B & C). There was no significant correlation between the ATL GM changes and ATL-connectivity after the control stimulation (intrinsic connectivity: r = - 0.23, p = 0.16, modulatory connectivity: r = 0.01, p = 0.49). Additionally, we examined the GM changes in the target site, ventrolateral ATL (MNI: -57, -15, -34) and found reduced GM, positively correlated with ATL-connectivity (Fig. S1). It is noted that there was no significant changes in occipital cortex following the Oz stimulation (control stimulation). No white matter changes were detected.

**Figure 3.**
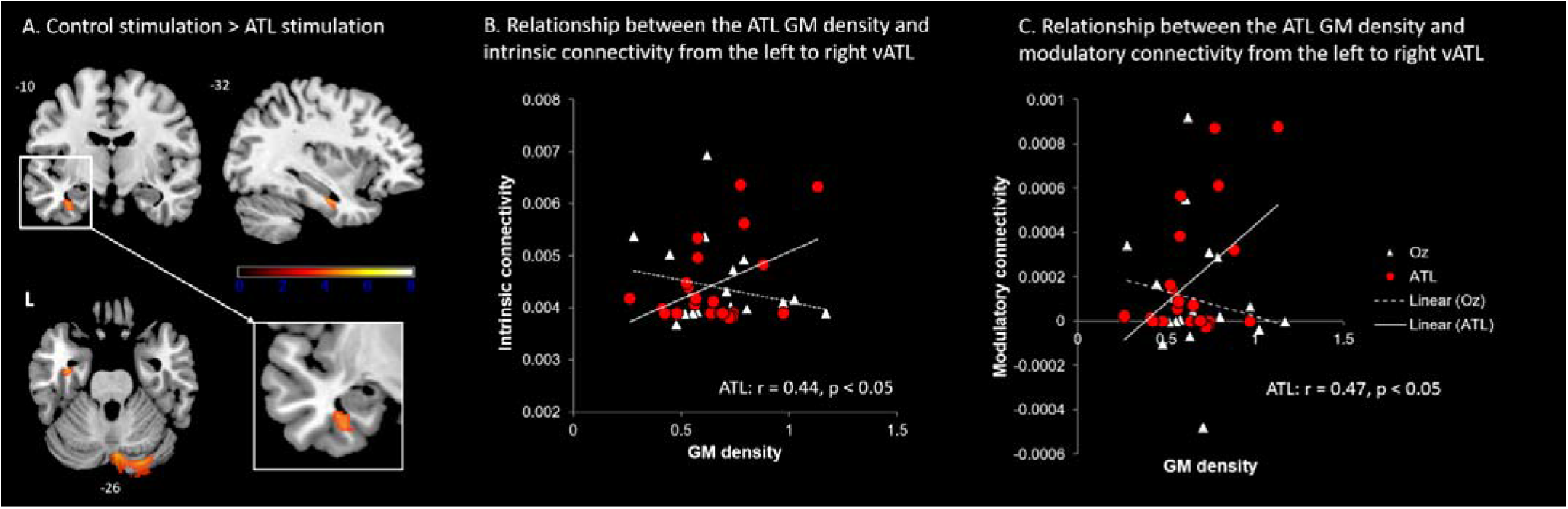
GM changes following cTBS over the left ATL compared to the control stimulation. A) Significantly decreased GM at the left ventral ATL and right cerebellum. B) The reduced ATL GM was positively correlated with intrinsic connectivity between the ATLs only after the ATL stimulation. C) The reduced ATL GM following the ATL stimulation was positively correlated with modulatory connectivity during semantic processing. Red circle represents the ATL stimulation. White diamond represents the control stimulation.

## 4. Discussion

Our results suggest that structural alterations in GM can occur very rapidly with one session of cTBS. Our results correspond to the time scale of TMS-induced structural plasticity in animal models (Funke and Benali, 2011) and learning-related morphological alterations in human (Sagi et al., 2012). Importantly, these local structural alterations were associated with the functional connectivity changes following cTBS. The local changes mirroring structural plasticity found in the current study support that TBS is able to produce powerful changes in regional synaptic activity in human adult brain (Siebner and Rothwell, 2003; Thomson et al., 2020). Our surprising results suggest a fast adaptive neural system supporting semantic processing reported previously (Jung and Lambon Ralph, 2016) and contribute to understanding of structural plasticity involved in clinical intervention with TMS.

Structural alterations induced by TMS is not well defined in human but animal studies demonstrated that rTMS/TBS induced immediate and prolonged structural plasticity in excitatory and inhibitory synapses (e.g., changes in size and number of related cells or receptors) in concomitant with functional plasticity (e.g., pre- and postsynaptic activity, neurotransmitter release) (Funke and Benali, 2011; Lenz et al., 2016). Recently, Thomson et al (2020) investigated the molecular mechanisms of TBS effects in living human neurons. They reported increase in the expression of plasticity genes following one-session of iTBS (600 pulses), compared to sham stimulation. Employing an *in vitro* human neuron-like model, they identified several gene expression changes supporting iTBS-induced plasticity. These studies provide strong evidence of immediate effect of TBS in structural plasticity. Here, we demonstrated that cTBS, an inhibitory TBS reduced the GM in the targeted region *in vivo*. VBM detects differences in local concentration or volume of GM and WM per voxel, changes in the classification of individual voxels, and potentially a combination of both (Ashburner and Friston, 2000). The underlying mechanism of GM changes includes axon sprouting, dendritic branching and synaptogenesis, neurogenesis, changes in glia number and morphology, and even angiogenesis (Zatorre et al., 2012). Although the GM change we observed might reflect alterations in cell genesis, the time scale of our study corresponds to fast adapting neuronal plasticity such as postsynaptic morphology changes (Funke and Benali, 2011; Lenz et al., 2016) and neuronal turnover (Trachtenberg et al., 2002), rather than slow mechanisms as neuronal or glia cell genesis (Kempermann et al., 1997).

Studies using TMS with functional imaging have demonstrated that unilateral rTMS/TBS leads to bilateral changes at a functional level, reflecting intrinsic connection between the stimulated region and functionally-connected remote homologous areas in the contralateral hemisphere in various cognitive function (Agnew et al., 2018; Andoh and Paus, 2011; Binney and Lambon Ralph, 2014; Hartwigsen et al., 2013; Jung and Lambon Ralph, 2016; Lee et al., 2003; O’Shea et al., 2007; Sack et al., 2005; Valchev et al., 2016). These studies suggest that rTMS/TBS can occur a compensatory short-term functional reorganization with increased contribution from the homologous area in the contralateral hemisphere after transient disruption delivered by inhibitory rTMS/TBS. Furthermore, May et al (2007) using VBM showed that 5 days iTBS over the left superior temporal gyrus induced increase and decrease GM changes at the bilateral superior temporal gyrus. Although we did not observe structural changes in the homologous right ATL following cTBS over the left ATL, our results showed that cTBS decreased GM and regional activity in the left ATL as well as changes in functional connectivity between the targeted ATL and homologous ATL in the contralateral hemisphere. Importantly, the GM alterations in the targeted region was associated with functional connectivity changes in the ATL-connectivity during semantic processing. Our findings suggest that TBS-induced structural neuroplasticity co-occurs with changes in functional processing and provide important insight about dynamic, fast-adaptive neural system (Siebner and Rothwell, 2003). Furthermore, these findings provide empirical evidence supporting theoretical framework that structural forms of plasticity serve an important function in processing information in neural systems when the flexible networks face novel information demands (Chambers et al., 2004).

In addition to the GM alterations in the left ATL following cTBS, we also observed GM reduction in the right cerebellum. May et al (2007) also reported GM increase in the cerebellum after stimulating the left superior temporal gyrus – the primary auditory cortex. Accumulating evidence suggest that the cerebellum is involved in language processing including word retrieval and generation and phonological and semantic processing (Marien et al., 2014) and has reciprocal connections with key language regions such as the left inferior frontal gyrus (IFG) and left lateral temporal gyrus (Booth et al., 2007). A recent study stimulated the right cerebellum with anodal transcranial direct stimulation (tDCS) and showed increased activation in the targeted cerebellum as well as left IFG, and posterior middle temporal gyrus (pMTG) during semantic predictive processing (D’Mello et al., 2017), which are the key regions in semantic cognition (Binder et al., 2009; Lambon Ralph et al., 2017). ATL has been considered as a transmodal hub in semantic representation interacting with semantic control regions including the IFG and pMTG (Lambon Ralph et al., 2017). Thus, the observed cerebellum GM decrease would be another functionally-connected remote changes within the semantic network following cTBS over the left ATL.

In neuroimaging studies, small to moderate sample size can contribute to false positive findings (Button et al., 2013). Thus, our results should be interpreted cautiously, taking into account the small sample size. We performed a power calculation in this study to estimate TBS effects in behaviour: 2 factorial within subject design with site (ATL vs. Oz) x task (semantic vs. control). The estimated interaction effect between the site and task was 0.36 with 23 participants. To achieve α=0.05, power=80% for the critical interaction between site and task, N≥18 are required. Although the power calculation showed that our sample size was sufficient to show the expected behavioural TBS effect, it remain unclear how many participants would be need to detect structural changes in VBM indices. With more subjects, we might see structural changes in the anterior commissure that we hypothesized as well as brain regions beyond the stimulation site. Further investigation will be needed to replicate these findings with larger sample size.

## Author contribution

J.J and M.A.L.R conceptualized and designed the experiment. J.J conducted the experiment and the data analyses. J.J and M.A.L.R wrote the manuscript.

## Conflict of interest

The authors declare no competing financial interests.

## Acknowledgments

This research was supported by a Beacon Anne McLaren Research Fellowship (University of Nottingham) to JJ and an Advanced ERC award (GAP: 670428 - BRAIN2MIND_NEUROCOMP) and MRC programme grant (MR/R023883/1) to MALR.

## Notes

### Competing Interest Statement

The authors have declared no competing interest.

