## Supplementary material for "Immediate impact of transcranial magnetic stimulation on brain structure: short-term neuroplasticity following one session of cTBS": SI

**Supplementary information**

**Supplementary Figure 1**

**Supplementary Figure 1**

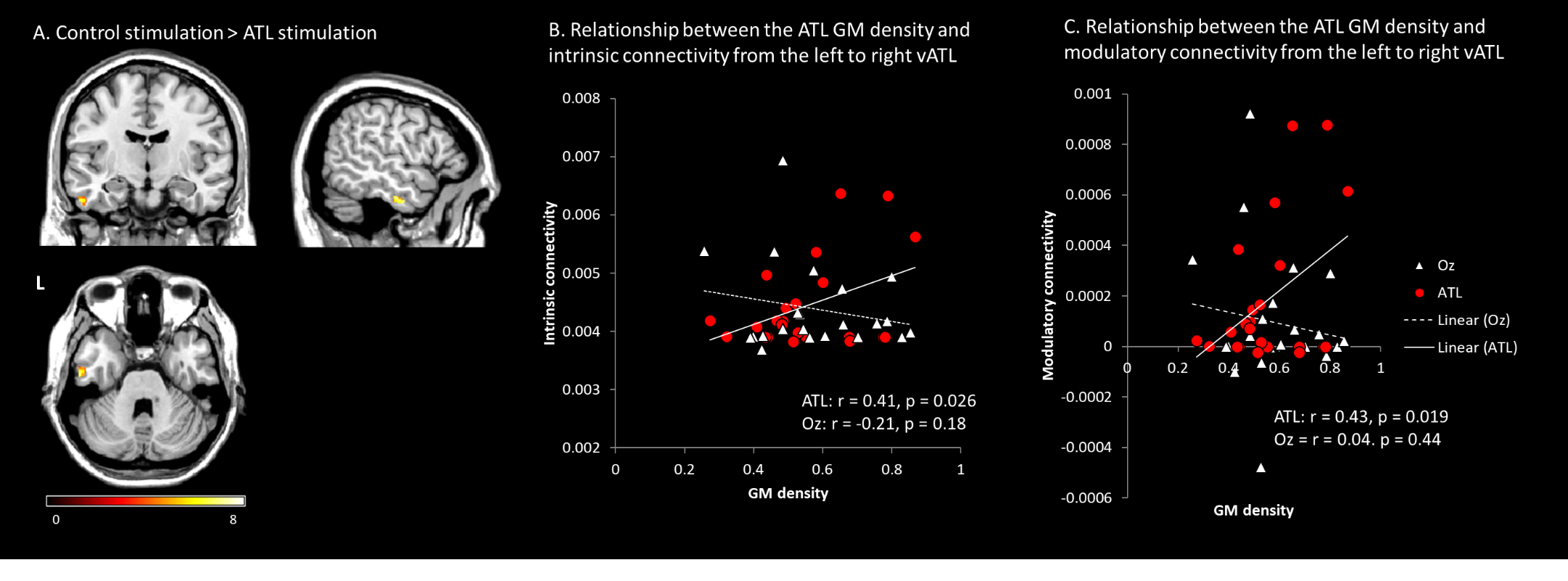

**Figure S1.** A) The results of VBM with the TMS target ROI (MNI: -57, -15, -34). B) Relationship between the ventrolateral ATL and the intrinsic connectivity from the left to right vATL. C) Relationship between the ventrolateral ATL and the modulatory connectivity from the left to right vATL. Red circles represent the ATL stimulation. White triangles represents the control (Oz) stimulation.
